## Supplemental text for "A computational method for predicting the most likely evolutionary trajectories in the step-wise accumulation of resistance mutations"

^1^R. Charlotte Eccleston

**This PDF file includes:**

Supplemental text

Figures S1 to S2

**Supplemental text**

**Analysis of mutation frequency data per country per region**

In what follows, we will describe in detail the frequency of the set of four *Pf*DHFR (N51I, C59R, S108N and I164L) and *Pv*DHFR (N50I, S58R, S117N, I173L) mutations found in each country of each region and attempt to infer evolutionary pathways. Due to the low number of isolates in some countries, the inferred pathways may not be reflective of the population and may change if the same analysis was carried out on a larger dataset. That is why we chose to carry out the analysis described in the main text on the larger combined datasets for the different regions. However, the below analysis indicates that the main trajectories followed in each region are relatively consistent within the countries of each region. The analysis below is carried out for all multi-country regions, but is not carried out for single-country regions (*Pf*DHFR: Middle Africa, Southern Asia and Melanesia; *Pv*DHFR: Central America, Eastern Asia and Melanesia), as this would be an identical analysis to the main text.

*Pf*DHFR

Western Africa:

In the main text, the main pathway to *Pf*DHFR quadruple mutation N51I,C59R,S108N,I164L in Western Africa was inferred from the isolate data to be S108N/C59R/N51I/I164L. Analysing the frequency of the mutations in each country included in the Western African isolates, the quadruple mutation is only found in Ghana. The most likely pathway to the quadruple mutation in Ghana is also S108N/C59R/N51I/I164L (S108N: 19/992; C59R,S108N: 160/992; N51I,C59R,S108N: 560/992; N51I,C59R,S108N,I164L: 1/992). The triple mutation N51I,C59R,S108N was the only triple mutation found in the isolates from Western Africa and was found in every country in the Western African dataset. We inferred the evolutionary trajectory to this triple mutation in each country where the quadruple mutation was not observed and found S108N/C59R/N51I was the most likely trajectory in Benin (S108N: 1/77; C59R,S108N: 1/77; N51I,C59R,S108N: 75/77), Cameroon (S108N: 1/239; C59R,S108N: 1/239; N51I,C59R,S108N: 234/239), Cote d’Ivoire (S108N: 1/70; C59R,S108N: 6/70; N51I,C59R,S108N: 35/70) and Guinea (S108N: 3/164; C59R,S108N: 9/164; N51I,C59R,S108N: 128/164). In Gambia, the most likely pathway was S108N/N51I/C59R (S108N: 2/250; N51I,S108N: 4/250; N51I,C59R,S108N: 216/250) and an alternative pathway S108N/C59R/N51I was also possible in this country (C59R,S108N: 3/250).

Single mutation S108N was absent from the isolates from Burkina Faso, Cape Verde, Gabon, Mali, Mauritania, and Nigeria but double and triple mutant combinations of the four mutations were observed. An alternative pathway to the triple mutation, C59R/S108N/N51I, was inferred in Guinea (C59R: 1/164; C59R,S108N: 9/164; N51I,C59R,S108N: 128/164) and Mali (C59R: 1/415; C59R,S108N: 23/415; N51I,C59R,S108N: 213/415). All single mutations N51I, C59R, S108N, I164L were absent from Burkina Faso, Cape Verde, Gabon, Mauritania and Nigeria and so complete pathways could not be inferred in these countries. This suggests either an insufficient number of samples were taken from these countries or evolution is at a later stage in these regions so only multiple mutations are observed. In Senegal, single mutation S108N (4/166), double mutation N51I,C59R (1/166) and triple mutation N51I,C59R,S108N (135/166) were observed, but all other combinations of the four mutations were absent. It is therefore not possible to infer a pathway for this country. The missing intermediate mutations suggests insufficient sampling of this region.

Eastern Africa:

The main pathway to the quadruple mutation inferred from Eastern African isolates in the main text was S108N/N51I/C59R/I164L with alternative pathway S108N/C59R/N51I/I164L. Considering the isolate data per country, the quadruple mutation N51I,C59R,S108N,I164L was found only in Kenya, however, all single mutations including S108N were absent from this country, making it difficult to infer the evolutionary pathway. However, by considering the frequency of double mutations we can infer alternative pathways. Double mutation N51I,S108N was observed in 14/118 isolates, triple mutation N51I,C59R,S108N was observed in 93/118 isolates and quadruple mutation N51I,C59R,S108N,I164L was found in 2/118 isolates. This suggests two likely pathways, N51I/S108N/C59R/I164L and S108N/N51I/C59R/I164L. Double mutation C59R,S108N is also observed in 7/118 Kenyan isolates. Therefore, we can infer two alternative pathways, C59R/S108N/N51I/I164L and S108N/C59R/N51I/I164L.

Triple mutation N51I,C59R,S108N is observed in all Eastern African countries included in the isolate data and so we will infer the evolutionary trajectories to this mutation for the remaining countries, where possible. In Malawi, the most likely pathway to this mutation is S108N/N51I/C59R (S108N: 2/366; N51I,S108N: 8/366; N51I,C59R,S108N: 352/366) with alternative pathway S108N/C59R/N51I (C59R,S108N: 4/366). Similarly, in Tanzania, the most likely pathway is S108N/N51I/C59R (S108N: 1/339; N51I,S108N: 28/339; N51I,C59R,S108N: 281/339) with alternative pathway S108N/C59R/N51I (C59R,S108N: 20/339).

All single mutations are missing in the remaining countries, so we infer possible pathways from the present double mutations. In Eritrea, double mutation N51I,S108N was found in 13/17 isolates and triple mutation N51I,C59R,S108N was found in 4/17 isolates, suggesting two possible pathways, N51I/S108N/C59R or S108N/N51I/C59R. Similarly, in Ethiopia, double mutation N51I,S108N was found in 4/25 isolates and triple mutation N51I,C59R,S108N was found in 21/25 isolates, suggesting possible pathways N51I/S108N/C59R and S108N/N51I/C59R.

In Madagascar and Uganda, only mutation N51I,C59R,S108N is present, suggesting insufficient sampling and making it impossible to infer pathways in these countries.

Southeastern Asia:

The main pathway inferred from the Southeastern Asian isolates was S108N/C59R/N51I/I164L with alternative pathway S108N/C59R/I164L/N51I also possible. Considering the countries in Southeastern Asia individually, the quadruple mutation is observed in all countries except Indonesia. However, single mutation S108N was only found in Cambodia, again suggesting either insufficient sampling or more advanced evolution in the remaining countries (the double mutation C59R,S108N and triple mutation N51I,C59R,S108N are high frequency in the countries where S108N in absent, except Indonesia). This makes it difficult to infer pathways for individual countries, with the exception of Cambodia. The pathway to the triple mutation in Cambodia is S108N/C59R/N51I/I164L (S108N: 3/1099; C59R,S108N: 71/1099; N51I,C59R,S108N: 521/1099; N51I,C59R,S108N,I164L: 492/1099). Although the single mutations are missing from the other regions, meaning we cannot infer the first step in the pathway, the presence of double and triple mutations enables us to infer what the likely first steps in the pathway are. In Laos, double mutation C59R,S108N was found in 47/126 isolates, suggesting the first step is either C59R or S108N. Triple mutation N51I,C59R,S108N was found in 70/126 isolates and quadruple mutation N51I,C59R,S108N,I164L was found in 1/126 isolates from Laos. This suggests two alternative pathways Laos S108N/C59R/N51I/164L or C59R/S108N/N51I/I164L. In Myanmar, double mutation C59R,S108N, triple mutations C59R,S108N,I164L and N51I,C59R,S108N and quadruple mutation N51I,C59R,S108N,I164L were found in 10/247, 32/247, 39/247 and 166/247 isolates, respectively. This suggests four alternative pathways C59R/S108N/I164L,N51I, S108N/C59R/I164L/N51I, C59R/S108N/N51I/I164L and S108N/C59R/N51I/I164L are all possible in this country. In Thailand, double mutations C59R,S108N and S108N,I164L, triple mutations C59R,S108N,I164L and N51I,C59R,S108N and quadruple mutation N51I,C59R,S108N,I164L were found in 47/932, 3/932, 52/932, 115/932 and 713/932. This suggests many possible pathways, the most likely being C59R/S108N/N51I/I164L, and S108N/C59R/N51I/I164L.

In Vietnam, single mutation I164L, was found in 1/245 isolates and double mutation S108N,I164L, triple mutation S108N,C59R,I164L and quadruple mutation N51I,C59R,S108N,I164L were found in 1/245, 5/245 and 47/245 isolates respectively, suggesting pathway I164L/S108N/C59R/N51I. The high frequency of double mutation C59R,S108N (11/245) and triple mutation N51I,C59R,S108N (175/245) suggests alternative pathways C59R/S108N/N51I/I164L or S108N/C59R/N51I/I164L are also possible.

South America:

In South America, evolution to triple mutation N51I,S108N,I164L was inferred to follow pathway S108N/N51I/I164L . In Peru, single mutation S108N was found in 14/23 isolates and triple mutation N51I,S108N,I164L was found in 4/23 isolates. The intermediates to the triple mutation, N51I,S108N or S108N,I164L are absent from this country, therefore there are two possible pathways to the triple mutation, S108N/N51I/I164L or S108N/I164L/N51I, although we cannot infer which one is more likely.

In Colombia, only single mutation S108N (12/23) and double mutation N51I,S108N (9/23) are observed, suggesting evolution occurs in the order S108N/N51I. In Brazil, double mutation N51I,S108N was found in 75% isolates, but single mutations N51I and S108N were absent.

*Pv*DHFR

In South America, evolution to triple mutation S58R,S117N,I173L was inferred to follow the main pathway S117N/S58R/I173L with alternative pathways S58R/S117N/I173L and S117N/I173L/S58R also possible. The triple mutation S58R,S117N,I173L was found in 2/85 isolates from Brazil. Single mutations S58R and S117N and double mutation S58R,S117N were found in 5/85, 7/85 and 24/85 Brazilian isolates, respectively. This suggests the most likely pathway to the triple mutation in Brazil is S117N/S85R/I173L, with an alternative pathway S58R/S117N/I173L. The triple mutation was not observed in any other South American countries. Double mutation S58R,S117N was observed in all South American countries except Guyana. Colombia (S117N: 24/34; S58R,S117N: 1/34) and Panama (S117N: 1/46; S58R,S117N: 1/46) appear to be following trajectory S117N/S58R to this trajectory, whilst Peru appears to follow S58R/S117N (S58R: 4/89; S58R,S117N: 48/89), with S117N/S58R possible but less likely (S117N: 2/89).

In Southeastern Asia, the triple mutation was not observed, but double mutation S58R,S117N was found in all countries except Malaysia. However, single mutations S58R and S117N were only found in isolates from Thailand. The inferred order of the mutations to this double mutation in Thailand was S58R/S117N (S58R: 4/160; S117N: 1/160; S58R,S117N: 6/160).

In Eastern Africa, evolution is only observed up to double mutation S58R,S117N. Eritrea (S117N: 1/13; S58R,S117N: 11/13) and Ethiopia (S117N: 24/53; S58R,S117N: 21/53) follow pathway S117N/S58R, whilst Sudan (S58R: 2/9; S117N: 1/9; S58R,S117N: 5/9) appears to follow the main pathway S58R/S117N with S117N/S58R also possible. In Madagascar 4/4 isolates contain S58R,S117N, therefore the low sampling in this country makes it impossible to infer the evolutionary order of the mutations. In Sudan S117N was found in 2/5 isolates, but the other combinations of mutations were absent.

The Southern Asian isolates also only contain evolution up to the double mutations S58R,S117N or N50I,S117N. In Afghanistan (S117N: 2/27; N50I,S117N: 4/27; S58R,S117N: 2/27) and Pakistan (S117N: 3/37; N50I,S117N: 2/37; S58R,S117N: 4/37), evolution appears to follow two possible pathways S117N/N50I and S117N/S58R. India appears to follow S58R/S117N (S58R: 3/48; S58R,S117N: 16/48) with alternative pathway S117N/S58R also possible (S117N: 1/48). In Bangladesh and Sri Lanka, all combinations of the four *Pv*DHFR mutations studied here are absent.

**The frequency of additional mutations in our isolate data**

*Pf*DHFR

Most of the mutations found in our isolate data were combinations of the four mutations N51I, C59R, S108N and I164L, as discussed above and in the main text. However, there were some additional mutations that were found in relatively high frequency in the isolate data. Triple mutation C59R,S108N,S306F was found in 1/1 isolates from Indonesia (Southeastern Asia) and 95/119 isolates from Papua New Guinea (Melanesia). Double mutation A16V,S108T was found in 3/1099 isolates from Cambodia (Southeastern Asia) and 1/23 isolates from Peru (South America) and single mutation A16V was found in 1/23 isolates from Colombia (South America). Single mutations N90Y and T130N were found in 1/415 isolates from Mali (Western Africa) and 1/52 isolates from Burkina Faso (Western Africa), respectively (See file ‘PfDHFR_isolate_frequencies.csv’).

*Pv*DHFR

There were many more diverse mutations in the isolates from *Pv*DHFR compared to *Pf*DHFR. Quadruple mutation F57I,S58R,T61M,N117T was found in 94/160 isolates from Thailand (Southeastern Asia) and 1/12 isolates from China (Southern Asia). Quadruple mutation F57L,S58R,T61M,N117T was found in 3/12 isolates from China (Southern Asia), 14/50 isolates from Malaysia (Southeastern Asia), 5/9 isolates from Indonesia (Southeastern Asia), 8/28 isolates from Myanmar (Southeastern Asia), 10/26 isolates from Papua New Guinea (Melanesia) and 26/160 isolates from Thailand (Southeastern Asia). Double mutation R58K,S117N was found in 11/85 isolates from Brazil (South America), 21/89 isolates from Peru (South America), 6/34 isolates from Colombia (South America), 6/53 isolates from Ethiopia (Ethiopia), 3/3 isolates from Guyana (South America), 1/50 isolates from Malaysia (Southeastern Asia) and 2/46 isolates from Panama (South America). Quadruple mutation F57L,I111L,N117T,I173F was found in 33/50 isolates from Malaysia (Southeastern Asia). Triple mutation R58K,S117N,I173L was found in 18/85 isolates from Brazil (South America), 1/38 isolates from Panama (South America), and 2/89 isolates from Peru (South America). Other mutations were found in lower frequencies, see file ‘PvDHFR_isolate_frequencies.csv’ for all mutation frequencies in the Supplementary data.

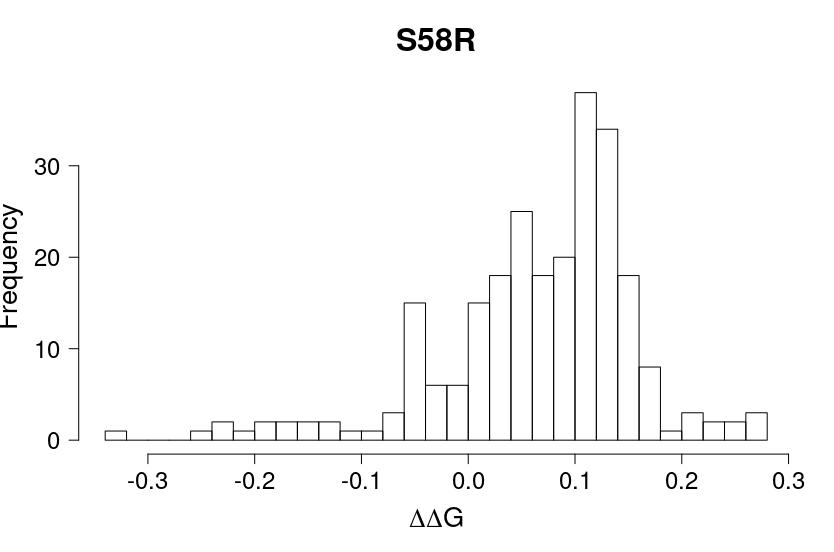

**Figure. S1:** Distribution of 250 Flex ddG predictions of *Pv*DHFR-pyrimethamine binding free energy change for single mutation S58R

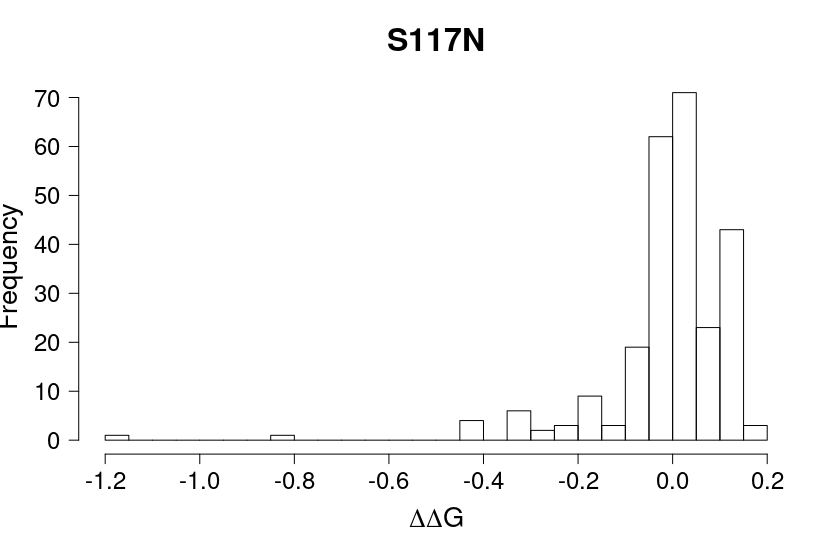

**Figure. S2.** Distribution of 250 Flex ddG predictions of *Pv*DHFR-pyrimethamine binding free energy change for single mutation S117N.
